# SOCS2 regulation of growth hormone signaling requires a canonical interaction with phosphotyrosine

**DOI:** 10.1101/2022.07.17.500381

**Authors:** Kunlun Li, Lizeth Meza-Guzman, Lachlan Whitehead, Evelyn Leong, Andrew Kueh, Warren S. Alexander, Nadia J. Kershaw, Jeffrey J. Babon, Karen Doggett, Sandra E. Nicholson

## Abstract

Suppressor Of Cytokine Signaling (SOCS) 2 is the critical negative regulator of growth hormone (GH) and prolactin signaling. Mice lacking SOCS2 display gigantism with increased body weight and length, and an enhanced response to GH treatment. Here we characterized mice carrying a germ-line R96C mutation within the SOCS2-SH2 domain, which disrupts the ability of SOCS2 to interact with tyrosine phosphorylated targets. *Socs2*^*R96C/R96C*^ mice displayed a similar increase in growth as previously observed in SOCS2 null (*Socs2*^*-/-*^) mice, with a proportional increase in body and organ weight, and bone length. Embryonic fibroblasts isolated from *Socs2*^*R96C/R96C*^ and *Socs2*^*-/-*^ mice also showed a comparable increase in phosphorylation of STAT5 following GH stimulation, indicating the critical role of phosphotyrosine binding in SOCS2 function.

## Introduction

Growth hormone (GH) is secreted by the anterior pituitary gland into the circulation, binding to a homo-dimeric GH receptor (GHR) on the surface of cells throughout the body to regulate growth, metabolism, and the immune response. In 1987, the GHR became the first cytokine/hematopoietin receptor superfamily member to be cloned, initiating a molecular investigation into GH signaling[1]. Despite this, it took many years to fully delineate mechanisms of ligand recruitment and signal propagation[2, 3]. GHR engagement promotes a conformational change in the pre-existing homodimeric receptor complex, activating the receptor-associated Janus kinase 2 (JAK2) tyrosine kinases[2, 3]. Activated JAK2 in turn phosphorylates tyrosine residues within the GHR cytoplasmic domain, recruiting Src homology 2 (SH2)-containing proteins such as the signal transducers and activators of transcription (STAT) proteins to the receptor complex[4]. Once STAT5b homodimers, the main mediators of GH signaling, are recruited to the receptor complex, they are phosphorylated by JAK2, triggering a conformational dimer change, translocation to the nucleus, and the transcription of genes involved in growth and development, such as insulin-like growth factor 1, *Igf1*[5-7]. Excessive GH production results in acromegaly in humans and is associated with severe pathological consequences[8], underscoring the need for cellular mechanisms that limit the GH response.

The SOCS protein family, consisting of CIS and SOCS1-7, functions to limit cytokine signaling, often in a negative feedback loop[9]. The important role of SOCS2 as a negative regulator of GH signaling was revealed by SOCS2-deficient mice, which displayed a 30-40% increase in growth compared to wild-type mice[10]. The gigantism was rescued by crossing SOCS2-deficient mice with *Ghrhr*^*lit/lit*^ (*little*) mice that lacked pituitary-derived GH secretion, with both *Ghrhr*^*lit/lit*^ and *Ghrhr*^*lit/lit*^ *Socs2*^*-/-*^ mice exhibiting comparable growth retardation[11].Similarly, SOCS2-null mice on a STAT5b-deficient background displayed a similar growth rate to wild-type mice, confirming SOCS2-deficient gigantism results from aberrant activation of GH signaling[12]. Notably, the increased weight of *Socs2*^*-/-*^ mice was not due to excess fat gain, but resulted from a proportional increase in organ size, including muscle and bone length. Consistent with this, SOCS2-deficient mice have longer femur, tibia, radius, and humerus bones[10], associated with increased bone mass, but not bone mineral density[13].

SOCS2 has a short 32-residue N-terminal region and two major functional domains: a central SH2 domain and a SOCS box. The SOCS2-SH2 domain is a substrate recognition module that binds to phosphorylated tyrosine residues 487 and 595 within the GHR cytoplastic domain, with SOCS2 binding proposed to inhibit cascade propagation via blocking access to other signaling intermediates[11]. The SOCS2-SH2 domain has also been reported to directly interact with JAK2[14]. The SOCS box provides binding sites for the adaptors Elongin B and C, and the Cullin 5 scaffold, which in turn, recruit RING-box protein (Rbx)2 to form an E3 ubiquitin ligase complex that ubiquitinates substrates bound to the SH2 domain[15]. There is an underlying assumption that SOCS2 ubiquitinates the GHR complex directing it to proteasomal degradation.

The SOCS2-SH2 domain structure consists of three β-strands flanked by two α-helices, and an additional “SOCS-specific” α-helix termed the Extended SH2 Subdomain (ESS)[16]. Interaction with phosphorylated targets occurs via a positively charged pocket (P0) that binds phosphorylated tyrosine (pTyr) and a hydrophobic patch that accommodates the third residue (+3) distal from the pTyr. Recently, we identified a non-canonical exosite on the SOCS2-SH2 domain that when occupied, enhances SH2 affinity for tyrosine phosphorylated targets[17].Multiple SOCS2 residues, including Arg96, coordinate binding to the pTyr residue[18]. A naturally occurring mutation of Arg96 to Cys in ovine SOCS2 is associated with inflammatory mastitis, and increased body size and milk production[19], with mutation of Arg96 disrupting SH2 binding to phosphopeptides[17, 19]. The characterisation of the R96C mutation in sheep was the first study to investigate the contribution of SH2:pTyr binding to SOCS2 function *in vivo*[19]. However, there are no comparative studies of SOCS2-deficient sheep to address whether SOCS2 function is completely reliant on its SH2 interaction with phosphotyrosine.

Here, we generated and characterized mice bearing the SOCS2-R96C mutation (*Socs2*^*R96C*^). Homozygous *Socs2*^*R96/R96C*^ mice displayed a 25-35% increase in weight compared to wild-type mice during puberty, and a similar increase in body weight to SOCS2-deficient mice. The gigantism resulted from a collective increase in weight of most visceral organs, associated with increased body and bone length. GH signaling was enhanced in *Socs2*^*R96C/R96C*^ fibroblasts, indicating that the augmented growth during development was a consequence of a dysregulated GH response. The *Socs2*^*R96C/R96C*^ mice displayed a similar phenotype to *Socs2*^*-/-*^ mice, highlighting a critical role for SOCS2-SH2 recognition of phosphotyrosine in canonical SOCS2 function.

## Results

### Mutation of R96 in SOCS2 does not disrupt SH2 domain integrity

SOCS2-SH2 binding to pTyr occurs through a cluster of residues consisting of Arg73, Ser75, Ser76, Thr83 and Arg96 in the SOCS2-SH2 domain[18] (**Figure 1A**), with all five residues conserved across different species (**Figure 1B**). Although Arg73 corresponds to the invariant arginine residue found in all SH2 domains[20-22], mutation of Arg73 to Lys in SOCS2 resulted in reduced the affinity for a phosphopeptide derived from GHR Y595 (pY595), but not loss of binding (**Supplementary Figure 1**).

**Figure 1.**
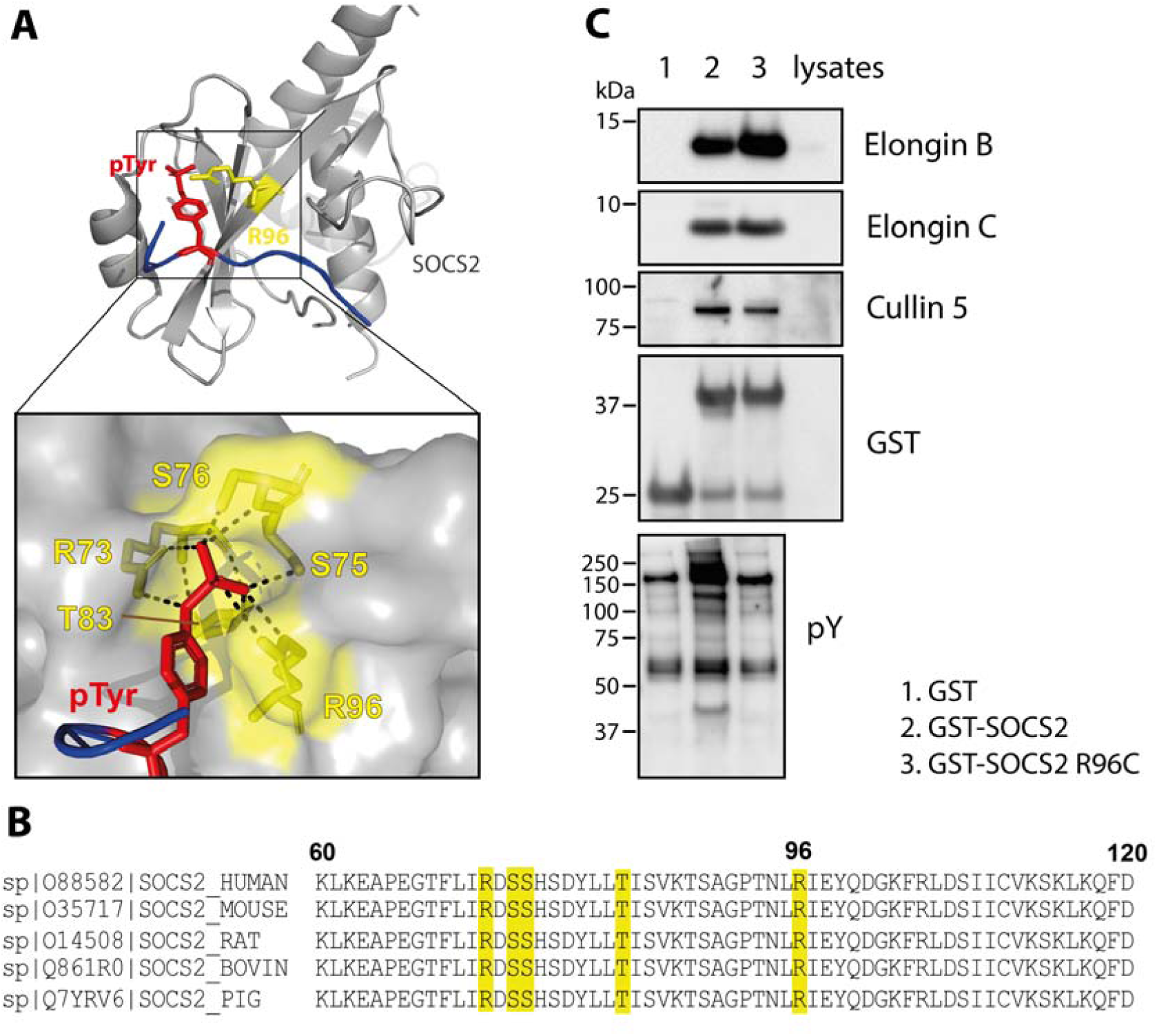
Arg96 within SOCS2 contributes to pTyr binding but not to SOCS box recruitment of the E3 ubiquitin ligase complex. (**A**) Crystal structure of SOCS2 bound to a phosphopeptide derived from Y426 in the erythropoietin receptor. Inset is a surface representation of the SOCS2-SH2 domain P0 pocket showing the hydrogen bond interactions (dashed lines) with phosphotyrosine (pY426 EPO-R peptide: red stick; Arg96: yellow stick). (Kung et al., Nat Comm 2019; PDB 6I4X). (**B**) Alignment of SOCS2 sequences from different species. Multiple sequences were obtained from UniProt database and aligned using UniProt alignment tool. Residues interacting with phosphotyrosine are highlighted in yellow. (**C**) Affinity enrichment of recombinant GST-SOCS2/BC and GST-SOCS2-R96C/BC proteins was performed with 293T cell lysates. GST-SOCS2 enrichment of interacting proteins was analyzed by immunoblotting with antibodies to Elongin B, Elongin C, Cullin5 and phosphotyrosine.

We had previously shown that mutation of Arg96 to Cys in the SOCS2-SH2 domain abrogated binding to a phosphopeptide derived from GHR Tyr595, without impacting on peptide binding to the SH2 exosite, confirming domain integrity[17]. To verify that the R96C mutation did not impact SOCS box function, GST-SOCS2 and GST-SOCS2-R96C, were purified in a trimeric complex with Elongins B and C, and used to affinity precipitate (AP) Cullin-5 from cell lysates. Elongin B and C were present at comparable levels in both GST-SOCS2 and GST-SOCS2-R96C complexes. GST-SOCS2-R96C efficiently enriched Cullin 5 to the same extent as GST-SOCS2, evidence that mutation of R96C within the SOCS2-SH2 domain did not disrupt the SOCS box E3 complex (**Figure 1C**). GST-SOCS2-R96C failed to enrich tyrosine phosphorylated proteins, confirming that pTyr binding was disrupted by the R96C mutation (**Figure 1C**).

### Gigantism in mice bearing a homozygous SOCS2 R96C mutation

To further investigate the impact of mutating Arg96 to Cys *in vivo*, we used CRISPR/Cas9 gene editing to generate a C57BL/6 mouse strain bearing the R96C mutation. Correct gene targeting was validated by next-generation sequencing (NGS), with routine genotyping performed by PCR (**Supplementary Figure 2**). Homozygous *Socs2*^*R96C/R96C*^ mice were viable and born at mendelian frequencies. *Socs2*^*R96C/R96C*^ mice were indistinguishable to *Socs2*^*R96C/+*^ and wild-type (WT) mice before weaning at three weeks of age, but subsequently developed at a faster rate (**Figure 2**). Analysis of immune cell populations from *Socs2*^*-/-*^ and *Socs2*^*R96C/R96C*^ mice showed increased numbers of splenic Natural killer (NK) cells (**Supplementary Figure 3**), consistent with previous observations[14]. By six weeks of age, both male and female *Socs2*^*R96C/R96C*^ mice weighed significantly more than *Socs2*^*R96C/+*^ or WT mice. The enhanced growth of *Socs2*^*R96C/R96C*^ mice was evident throughout puberty, with growth plateauing at week 8, and maintained in adult mice (**Figure 2C-F**). At 8-weeks of age, body weights of *Socs2*^*R96C/R96C*^ mice were comparable to *Socs2*^*-/-*^ mice, with both significantly heavier than WT mice (30.7% and 29.1% in male, 25.4% and 27.5% in female, respectively), suggesting that the weight difference in *Socs2*^*R96C/R96C*^ mice phenocopied *Socs2*^*-/-*^ mice (**Figure 2B**). At 12-weeks of age, *Socs2*^*R96C/R96C*^ male and female mice were visually different to WT mice (**Figure 2A** and not shown), with *Socs2*^*R96C/R96C*^ male mice attaining a 31.5% weight increase over WT males (*Socs2*^*R96C/R96C*^: 37.1 ± 2.42g, WT: 28.2 ± 1.39g) (**Figure 2C, D**). *Socs2*^*R96C/R96C*^ females typically attained the weight of WT male mice and displayed a 26.5% weight increase over WT females at 12-weeks of age (*Socs2*^*R96C/R96C*^: 27.2 ± 1.53g, WT: 21.5 ± 0.98g) (**Figure 2E, F**). Male and female *Socs2*^*R96C/+*^ mice exhibited similar development and adult body weights to WT mice (Figure 2C-F). The increased size of *Socs2*^*R96C/R96C*^ *mice* was reflected in a proportional increase in the weight of individual organs, with the majority of organs (excluding brain) showing increased weight compared to organs from WT mice. Pancreas and testes in males did not show a statistical difference. Carcass weight and body length were also increased in *Socs2*^*R96C/R96C*^ mice (**Figure 2G** and not shown).

**Figure 2.**
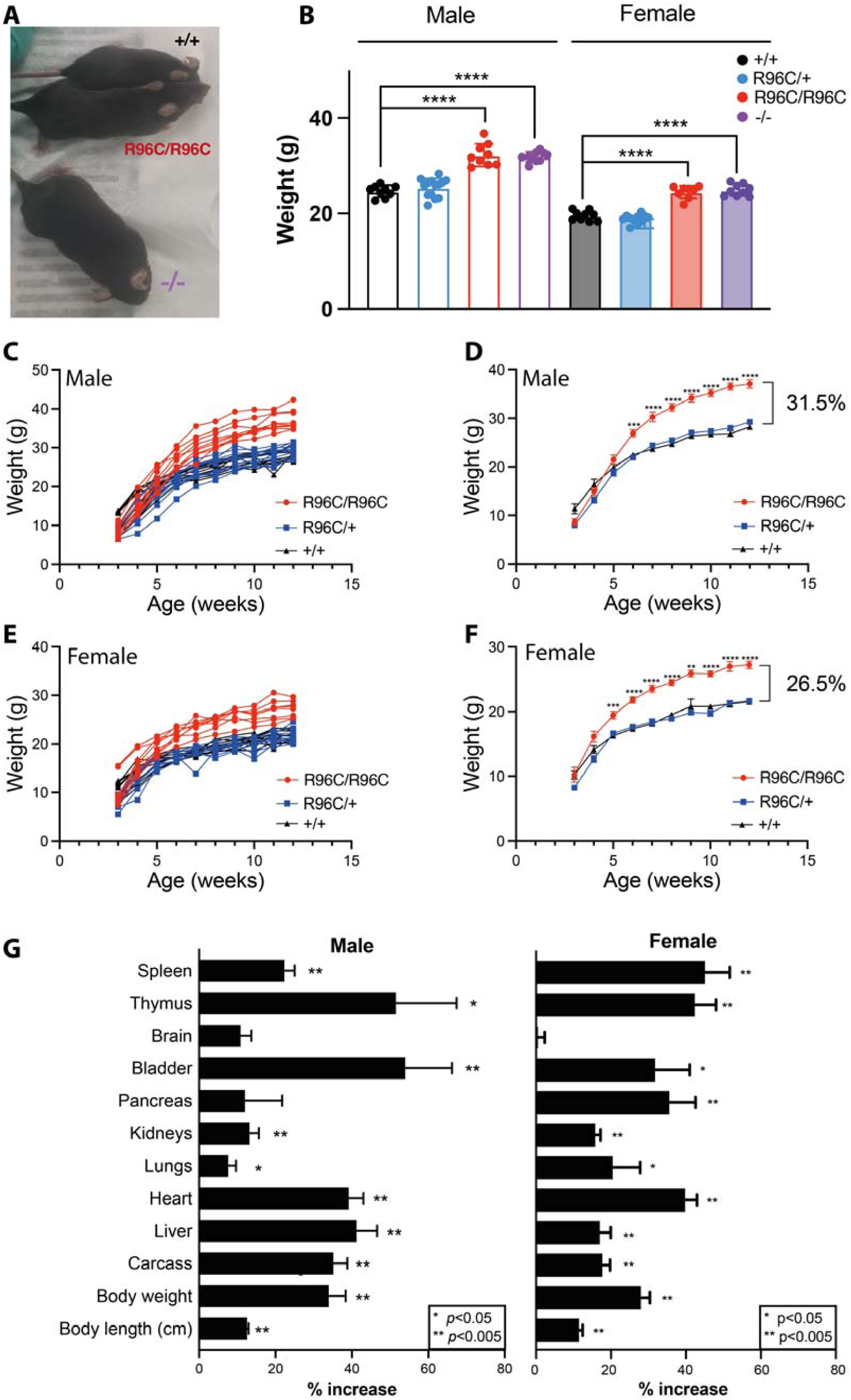
Homozygous *Socs2*^*R96C/R96C*^ mice display enhanced growth consistent with dysregulated growth hormone signaling during puberty. (**A**) 13-week-old wild-type (+/+), *Socs2*^*R96C/R96C*^ and *Socs2*^*-/-*^ male mice. (**B**) Body weights of 8-week-old wild-type (+/+), *Socs2*^*R96C/R96C*^ and *Socs2*^*-/-*^ mice. Each dot represents an individual mouse. (**C**) Growth curves from male littermates: *Socs2*^*R96C/R96C*^ (n=9), *Socs2*^*R96C/+*^ (n=14), *Socs2*^*+/+*^ (n=9). **(D)** Data from panel C as mean ± S.E.M. **(E)** Growth curves from female littermates: *Socs2*^*R96C/R96C*^ (n=8), *Socs2*^*R96C/+*^ (n=11), *Socs2*^*+/+*^ (n=9). **(F)** Data from panel E as mean ± S.E.M. Mice were weighed weekly from 3-weeks of age. **(H)** Body length and carcass and organ weights were measured for 13-week-old male *Socs2*^*R96C/R96C*^ (n=7) and *Socs2*^*+/+*^ (n=7), and female *Socs2*^*R96C/R96C*^ (n=8) and *Socs2*^*+/+*^ (n=6) mice. Data are shown as mean ± S.E.M. and expressed as a % of wild-type averages. Data were analyzed using an unpaired *t*-test. *p<0.05, **p<0.005, ***p<0.0005, ****p<0.00005.

**Figure 3.**
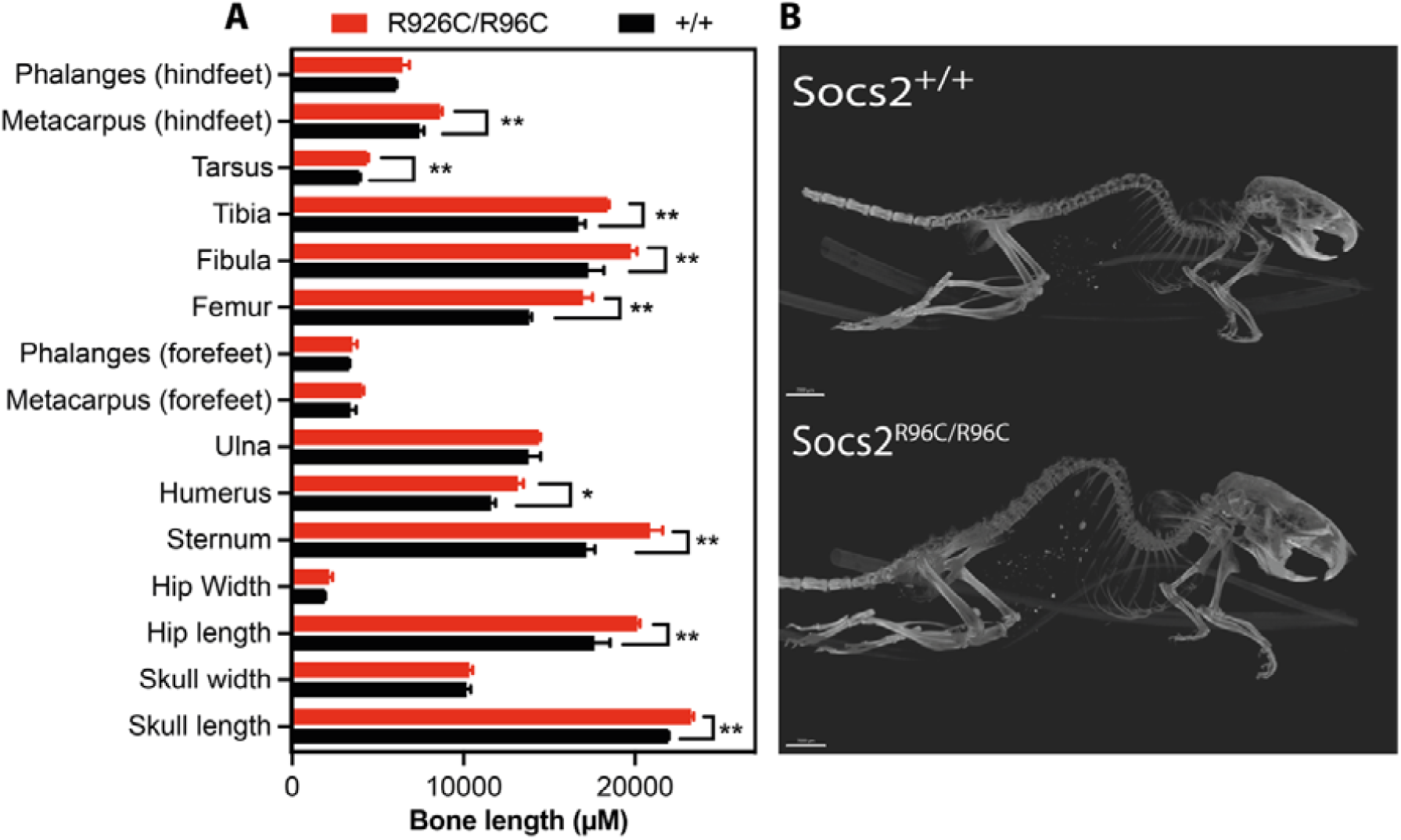
Homozygous Socs2-R96C mice display enhanced bone growth. 14-week-old *Socs2*^*R96C/R96C*^ and *Socs2*^*+/+*^ male mice were euthanized and whole animals analysed by micro computed tomography (micro-CT). Skeleton X-ray imaging in 3D was collected using a Bruker Skyscan 1276 Micro-CT. The X-ray projection images were reconstructed into 3D volumes using Bruker’s NRecon software, skeletons were visualized, and individual bone lengths measured in Imaris 8.7. (A) Bone lengths were shown in mean ± S.D. (n=3) and analysed using *t*-tests *p<0.05, **p<0.005. (B) Example micro-CT images.

Micro computed tomography (CT) X-ray imaging of skeletons from 13-week-old male WT and *Socs2*^*R96C/R96C*^ mice revealed an enlarged skeleton and increased bone length of skull, hip, sternum, humerus, femur, fibula, tibia, tarsus and metacarpus (hindfeet) in *Socs2*^*R96C/R96C*^ mice. Histology of skin from 13-week *Socs2*^*R96/R96C*^ male mice revealed dermal thickening and increased collagen production (**Supplementary Figure 4)**, consistent with skin pathology in *Socs2*^*-/-*^ mice[10].

**Figure 4.**
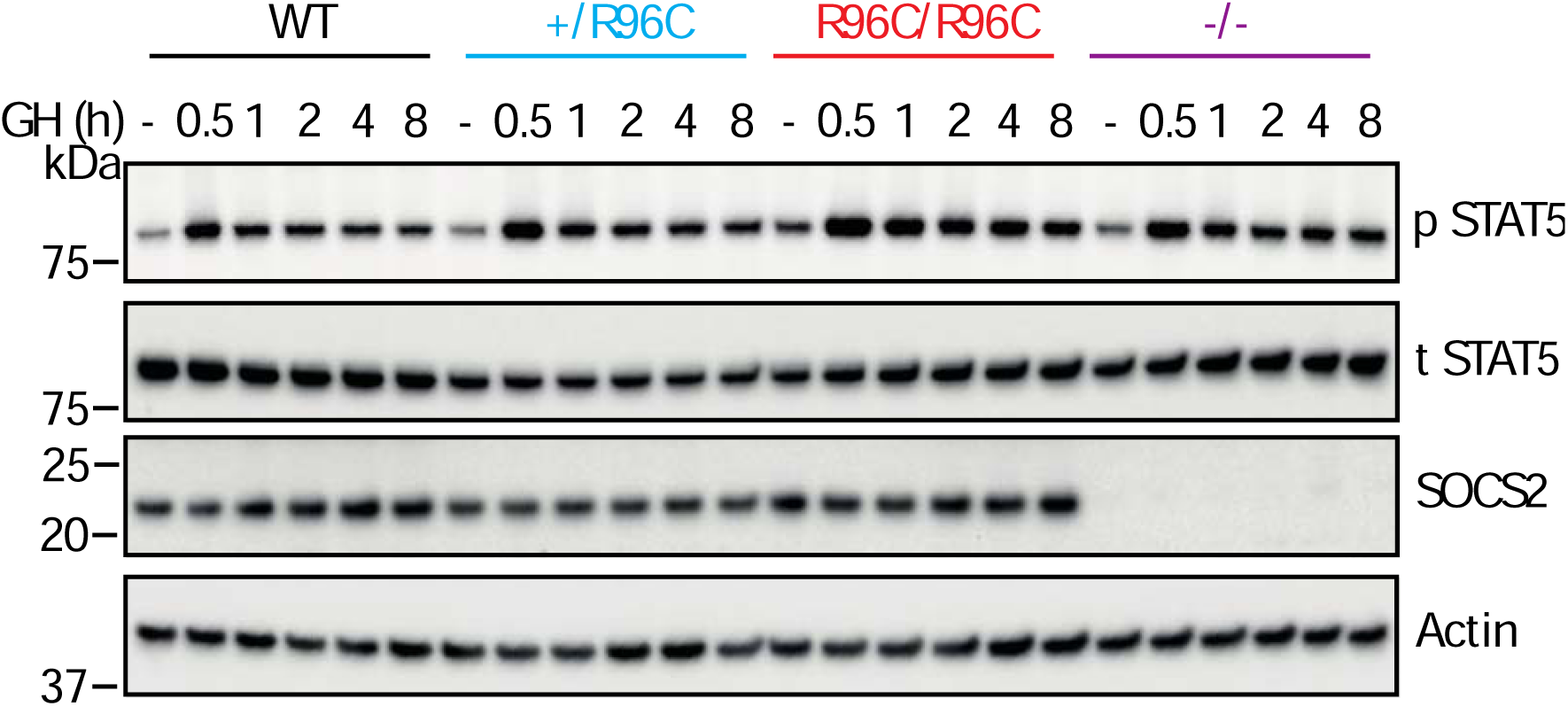
*Socs2*^*R96C/R96C*^ and *Socs2*^*-/-*^ MEFs display prolonged growth hormone signaling. *Socs2*^*+/+*^, *Socs2*^*R96C/+*^, *Socs2*^*R96C/R96C*^ and *Socs2*^*-/-*^ MEFs were treated with and without (-) 50 ng/mL of GH, lysed and analysed by immunoblotting with antibodies to the indicated proteins. P: phosphorylated, T: total. Representative of three independent experiments. Related to **Supplementary Figure 5**.

### Loss of SOCS2-SH2:pTyr binding results in enhanced GH signaling

To further investigate the impact of the SOCS2-R96C mutation on GH signaling, mouse embryonic fibroblasts (MEFs) were generated from *Socs2*^*+/+*^, *Socs2*^*R96C/+*^, *Socs2*^*R96C/R96C*^, and *Socs2*^*-/-*^ day-13 embryos. GH stimulation resulted in robust tyrosine phosphorylation of STAT5 (pSTAT5; **Figure 4 and Supplementary Figure 5**), which peaked at 0.5 h GH treatment in MEFs of all genotypes. The intensity of the pSTAT5 response was comparable in *Socs2*^*R96C/R96C*^ and *Socs2*^*-/-*^ MEFs, and consistently increased at 1-8 h GH treatment, relative to *Socs2*^*R96C/+*^and *Socs2*^*+/+*^ MEFs. *Socs2*^*R96C/+*^ MEFs displayed an intermediate level of pSTAT5 compared to WT and *Socs2*^*R96C/R96C*^ MEFs. Total STAT5 levels were comparable in all genotypes. SOCS2 was present at baseline, reduced in *Socs2*^*R96C/+*^ MEFs and expressed at comparable levels in WT and *Socs2*^*R96C/R96C*^ MEFs. There was a modest upregulation of SOCS2 at 8 h. These data indicate that loss of phosphotyrosine binding is sufficient to abrogate SOCS2 regulation of GH signaling.

## Discussion

We had previously shown that the SOCS2-R96C mutation did not impact on F3 peptide binding to the SOCS2 exosite[17]. In this study, we confirmed the R96C mutation did not alter the ability of SOCS2 to recruit the SOCS box-associated-E3 ubiquitin ligase complex. Taken together, this confirmed the R96C mutation did not compromise domain integrity, enabling us to interrogate the role of SOCS2-SH2:pTyr binding in mice. *Socs2*^*R96C/R96C*^ mice displayed increased growth during puberty, attaining a ∼25% and ∼30% increase in weight (females and males) compared to heterozygous littermates. The enhancement in body size and weight was comparable to SOCS2 null mice. In addition, primary embryonic fibroblasts derived from *Socs2*^*R96C/R96C*^ mice displayed prolonged GH-induced signaling compared to WT cells, consistent with loss of a negative regulator (SOCS2), and further evidence that loss of phosphotyrosine binding was sufficient to fully disrupt SOCS2 function.

The structure of SOCS2 in complex with the GHR pY595 peptide indicated that SOCS2 has the capacity to not only engage phosphopeptide in the conventional manner, but can also interact with a second GHR pY595 peptide orientated in an anti-parallel direction. This model provides a potential mechanism for SOCS2 interaction with both subunits in the dimerized GHR[18]. In addition to pTyr binding, the SOCS2-SH2 domain can also bind to a non-phosphorylated peptide via the exosite, providing an alternative site for SH2 interaction with protein targets[17]. These two non-canonical interaction sites within the SOCS2-SH2 domain suggest there may be additional protein interactions and regulatory mechanisms that impact on SOCS2 function. While our results confirm the critical role of SOCS2 in regulating growth hormone signaling, the genetic relationship to mastitis in sheep (a bacterial infection of the udder)[19], indicates SOCS2 also has an important role in regulating the immune response to bacterial infection, most likely related to its upregulation by LPS and IFNγ, and inhibition of NFkB signaling[23, 24]. This suggests that SOCS2 regulates proteins independently of GHR signaling complexes and this may occur either through classic SH2:pTyr interactions or via non-canonical SH2 interactions.

*SOCS2* has been associated with reduced growth in children and diverse disease pathologies. Given that mouse and human *SOCS2* share 94% sequence similarity, they are likely to have a conserved role as a critical negative regulator of GH signaling[12, 17]. This is illustrated by correlation of a *SOCS2* polymorphism with a positive response to GH therapy during puberty, and increased adult height of children with GH deficiency and Turner syndrome after long-term rhGH treatment[25]. Thus, inhibition or reduction of SOCS2 has the potential to enhance the efficiency of long-term rhGH therapy. *SOCS2* mRNA levels were decreased in osteoarthritis patients[26], and a polymorphism in the *SOCS2* gene correlated with susceptibility to type 2 diabetes in a Japanese population[27]. Reduced SOCS2 protein expression is associated with a poorer outcome in diverse human cancers, such as colorectal cancer[28], breast carcinoma[29] and hepatocellular carcinoma[30], suggesting SOCS2 may have some utility as a prognostic biomarker.

SOCS2 is also an important regulator in the immune system[31]. DC-mediated T cell priming and adaptive anti-tumoral immunity is enhanced in SOCS2-deficient mice, resulting in reduced tumor burden[32]. Loss of SOCS2 in CD4^+^ T cells results in increased T helper (Th) 2 cell differentiation and allergic inflammation[33], while *Socs2*^*-/-*^ CD4^+^ cells also show reduced Treg differentiation *in vivo* and *in vitro* under TGF*β* stimulation[34]. *Socs2*^*-/-*^ mice display increased numbers of NK cells (consistent with our observations) and upregulated JAK2 and STAT5 activation in response to IL-15[14].

The canonical role of SOCS2 in regulating growth hormone signaling cannot account for the diverse role of SOCS2 in immunity and inflammatory diseases. The *Socs2*^*R96C/R96C*^ mice provide an accessible model to investigate candidate pTyr targets for the SOCS2-SH2 domain *in vivo*, which will enable us to dissect the different pathways regulated by SOCS2, as well as the identification of novel SOCS2 targets.

## Data availability

All supporting data are included within the main article and supplementary file.

## Acknowledgements

We thank Dr Andrew Brooks (The University of Queensland Diamantina Institute) for providing human growth hormone. J.J.B was supported by an Australian Government National Health and Medical Research Council (NHMRC) Research Fellowship (1121755).K.L. was supported by a Melbourne Research Scholarship (University of Melbourne). This work was supported under a collaborative research project with Servier, and was supported in part through Victorian State Government Operational Infrastructure Support and the Australian Government NHMRC Independent Research Institutes Infrastructure Support Scheme (IRIISS).

## Methods

### Protein purification

pGEX4T constructs for expression of GST-human SOCS2 (residues 32–198), SOCS2-R73K (residues 32–198) and SOCS2-R96C (residues 32–198), and pACYCDuet constructs for expression of human ElonginB (residues 1-118) and ElonginC (residues 17-112), have been described previously[17]. The GST-SOCS2/BC trimeric complex was produced by co-expression in *E. coli* and purified by affinity chromatography as described previously[17]. The R73K mutation was introduced using the QuikChange Lightning Site-Directed Mutagenesis Kit (Agilent Technologies). The primer for mutagenesis (AATGCGAGCTATCTTTAATCAAGAAAGTTCCTTCTGGTGCC) was from Integrated DNA Technologies (Singapore).

### GST affinity precipitation

Human Embryonic Kidney cells (HEK293T) cells were pre-treated with 10 µM MG132 for 4 h and sodium pervanadate for the final 30 min[35]. 2 × 10^7^ cells were lysed in 1 mL NP-40 lysis buffer (1% v/v NP-40, 50 mM HEPES, pH 7.4, 150 mM NaCl, 1 mM EDTA, 10% glycerol, 1 mM PMSF, 1 mM Na_3_VO_4_, 1 mM NaF and protease inhibitors (Complete Cocktail tablets, Roche). Cell lysates were clarified by centrifugation at 13,000 g for 15 min at 4 °C. Supernatant was pre-cleared with glutathione-sepharose resin (GE Healthcare). 5 µg GST or 10 µg GST-SOCS2 protein was then added to cell lysates and incubate for 2.5 h at 4 °C. 30 µl of 50% Glutathione Sepharose resin (GE Healthcare) was then added and incubate for 60 min at 4 °C. Glutathione Sepharose resins were then washed with 3 × 1 mL of NP-40 lysis buffer and boiled with 20 μl SDS sample buffer.

### Surface plasmon resonance (SPR)

The affinity of SOCS2 and SOCS2-R73K protein for a phosphopeptide derived from the GHR receptor was measured by competitive SPR, as described previously[17]. Briefly, biotin-GSGS-GHR pY595 peptide was immobilised to a Streptavidin-coated SA chip and 100 nM of SOCS2 proteins pre-incubated with titrations of GHR pY595 competitor peptide (10 µM, 3.3 µM, 1.1 µM, 0.3 µM, 0.1 µM), were flowed over the chip. Peptides were purchased from Genscript.

### Mice

Mice carrying a germline mutation of Arg96 to Cys in SOCS2 (Socs2^R96C^) were generated on a C57BL/6LJ background by the MAGEC laboratory at the Walter & Eliza Hall Institute, as previously described[36]. 20 ng/µL Cas9 mRNA, 10 ng/µL single guide RNA (CAGCTGGACCGACTAACCTG), and 40 ng/µL oligo donor template (TCGCATTCAGACTACCTACTAACTATATCCGTTAAGACGTCAGCTGGACCGACTAAtCTatGtATTGAGTACCAAGATGGGAAATTCAGATTGGATTCTATCATATGTGTCAAGTCCAAGCTT) were injected into fertilized one-cell stage embryos. Additional silent changes were incorporated into the donor template to enable genotyping by PCR. Founder mice were analyzed by next-generation sequencing (NGS) to confirm the correct sequence change. Mice carrying the mutation were backcrossed to C57BL/6J mice for three generations to eliminate potential off-target events, and next-generation sequencing repeated. Heterozygous *Socs2*^*R96C/+*^ mice were intercrossed to generate a homozygous *Socs2*^*R96C/R96C*^ line. Mice were routinely genotyped using genomic DNA extracted from ear biopsies with the Direct PCR Lysis tail reagent (Viagen) and 5□mg/mL proteinase K (Worthington), according to the manufacturer’s instructions. Sequencing primers are shown in **Supplementary Table 1**. *Socs2*^*-/-*^ mice have been described previously[10], and both *Socs2*^*-/-*^ and wild-type (WT) C57BL/6 mice were bred and housed in the same room as *Socs2*^*R96C*^ mice. All experiments were approved by the WEHI Animal Ethics Committee and performed in accordance with the NHMRC Australian code for the care and use of animals for scientific purposes.

### Micro-computed tomography (micro-CT)

14-week-old *Socs2*^*R96C/R96C*^ and WT mice (n=3) were euthanized, and skeletal bones analysed in intact carcasses by skeleton X-ray imaging in 3D using a Bruker Skyscan 1276 Micro-CT. X-ray images were reconstructed into 3D volumes using Bruker’s NRecon software, skeletons were visualised, and length of individual bones measured in Imaris 8.7.

### Skin histology

Dorsal skin sections were taken from 10-week-old male *Socs2*^*R96C/R96C*^, *Socs2*^*-/-*^ and WT mice (n=3), fixed with 10% buffered formalin, embedded in paraffin and stained with Van Giessen stain using standard techniques. Images were captured using the Panoramic Scan II (3D HISTECH Ltd.).

### Immunophenotyping

Spleens, axial and inguinal lymph nodes, and cardiac bleeds were collected from 8-12-week-old mice. Single cell suspensions from spleens and lymph nodes were acquired by mashing organs through a 70 µM cell strainer with FACS buffer (phosphate-buffered saline (PBS) + 2% FCS). Cardiac bleeds were collected in K3-EDTA tubes, and whole blood separated by density gradient, underlaying Histopaque®-1077 (Sigma-Aldrich) and centrifuging at room temperature for 15 min at 400 x g with low acceleration and brake. The opaque interface was collected and washed twice with FACS buffer. Spleen and lymph node samples were resuspended in 5 mL of red cell removal buffer (156 mM NH_4_Cl, 11.9 mM NaHCO_3_ and 0.097 mM EDTA) for 5 min at room temperature and washed twice with PBS.

Single cell suspensions were resuspended in FACS buffer and 20 µL removed to determine absolute cell numbers, using 123count eBeads (Invitrogen™; 10,000 beads/tube) and propidium iodine (1 µg/mL). The remaining cells were incubated with Zombie UV (1:1000 in PBS) for 15 min at room temperature, washed twice with FACS buffer, FC blocked (FcR Blocking Reagent, mouse, Miltenyi;1:100 in FACS buffer) for 10 min at 4°C in the dark, and stained for 30 min at 4°C with the relevant antibody cocktail (**Table 1**). Cells were washed twice with PBS and fixed with 100 µL BD Cytofix™ Fixation Buffer for 30 min at 4°C. Fluorescent labelling and fixation was performed in the dark. Finally, cells were washed twice with PBS and analysed using a BD FACSymphony™ cell analyser. Data analysis was performed with FlowJo software v10.

**Table 1.** Antibodies used in flow cytometric analysis.

| Antigen | Fluorophore | Dilution | Clone | Company | Catalogue number |
| --- | --- | --- | --- | --- | --- |
| NK1.1 | FITC | 1:100 | PK136 | BD* | 553164 |
| TCR | percp-cy5.5 | 1:400 | H57-597 | BD | 560657 |
| CD3e | percp-cy5.5 | 1:400 | 145-2C11 | BD | 551163 |
| CD4 | BV421 | 1:1000 | RM4-5 | Biolegend | 100546 |
| CD8a | BV650 | 1:800 | - | BD | 563234 |
| CD49b | APC | 1:200 | DX5 | BD | 560628 |
| CD19 | AF700 | 1:200 | 1D3 | BD | 557958 |
| CD45 | APC-CY7 | 1:200 | 30-F11 | BD | 557569 |
| NKp46 | PE/CY7 | 1:100 | 29a1.4 | eBioscience | 25-3351-82 |

\* BD Biosciences

### Isolation of primary mouse embryonic fibroblasts (MEFs) and GH stimulation

Male and female *Socs2*^*R96C/+*^ mice were set-up in timed matings, and e13 embryos harvested to generate *Socs2*^*+/+*^, *Socs2*^*R96C/+*^ and *Socs2*^*R96C/R96C*^ MEFs. *Socs2*^*-/-*^ MEFs were generated from *Socs2*^*-/-*^ matings. Heads and fetal livers were removed, and embryos mechanically disaggregated and digested using trypsin, before culturing in tissue culture plates coated in 0.1% gelatin/PBS. MEFs were cultured in DMEM (Thermo) supplemented with 100□/mL penicillin, 0.1□g/ml streptomycin and 10% FBS (Thermo). at 37°C in a humidified incubator with 10% CO_2_. MEFs at passage 5 were treated with 50 ng/mL of human GH for 0.5 to 8 h, lysed in NP-40 lysis buffer, and analysed by immunoblotting. Human GH was kindly provided by Dr Andrew Brooks.

### Immunoblotting

Cell lysates were separated by sodium dodecyl sulphate-polyacrylamide gel electrophoresis (SDS-PAGE) and electrophoretically transferred to nitrocellulose membranes (Amersham). Membranes were blocked overnight in 5% w/v BSA and incubated with primary antibody for 2 h. Anti-GST-HRP (RPN1236, 1:10000) was from Sigma Aldrich. Anti-phosphotyrosine antibody (4G10) was obtained from Millipore. Anti-Elongin C antibody (610761, 1:3000) was from BD Biosciences. Anti-p44/42 (9102S, 1:2000), anti-STAT5 (94205; 1:3000) and anti-phospho-STAT5 (9359; 1:2000) were purchased from Cell Signaling. Anti-actin-HRP antibody (C4) was obtained from Santa Cruz (sc-47778 HRP; 1:1000). Anti-Cullin 5(ab184177), anti-Elongin B antibody (ab154854, 1:2000) and anti-SOCS2 antibody (ab109245, 1:1000) were purchased from Abcam. Antibody binding was visualized with peroxidase-conjugated sheep anti-rabbit immunoglobulin (Southern Biotech; 4010-05; 1:15000), or sheep anti-mouse immunoglobulin (GE Healthcare; NA931-1ML; 1:10000) and the enhanced chemiluminescence (ECL) system (Amersham or Millipore).

## Statistical analysis

Statistical analyses were performed with Prism 9 software. All statistical parameters including the exact value of n, the statistical test, error bars and significance are reported in associated figure legends.

## Notes

### Competing Interest Statement

The authors have declared no competing interest.

