## Supplementary for "SOCS2 regulation of growth hormone signaling requires a canonical interaction with phosphotyrosine"

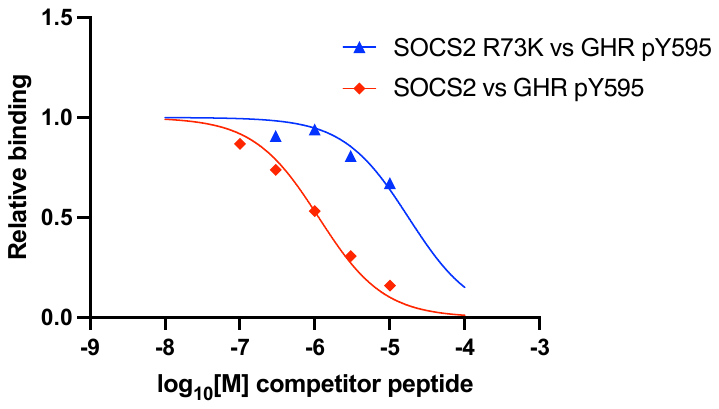

**Supplementary Figure 1. Mutation of Arg73 to Lys in the SOCS2-SH2 domain results in reduced binding to phosphopeptide.** A competitive surface plasmon resonance (SPR) assay was used to assess the impact of the R73K mutation. SOCS2 bound to a phosphopeptide derived from GHR pY595 with an IC_50_ 1.1 ± 0.05 μM. The SOCS2-R73K mutation reduced binding to GHR pY595, IC_50_ 26.7 ± 1.0 μM. IC_50_ values are mean ± S.D., derived from three independent experiments.

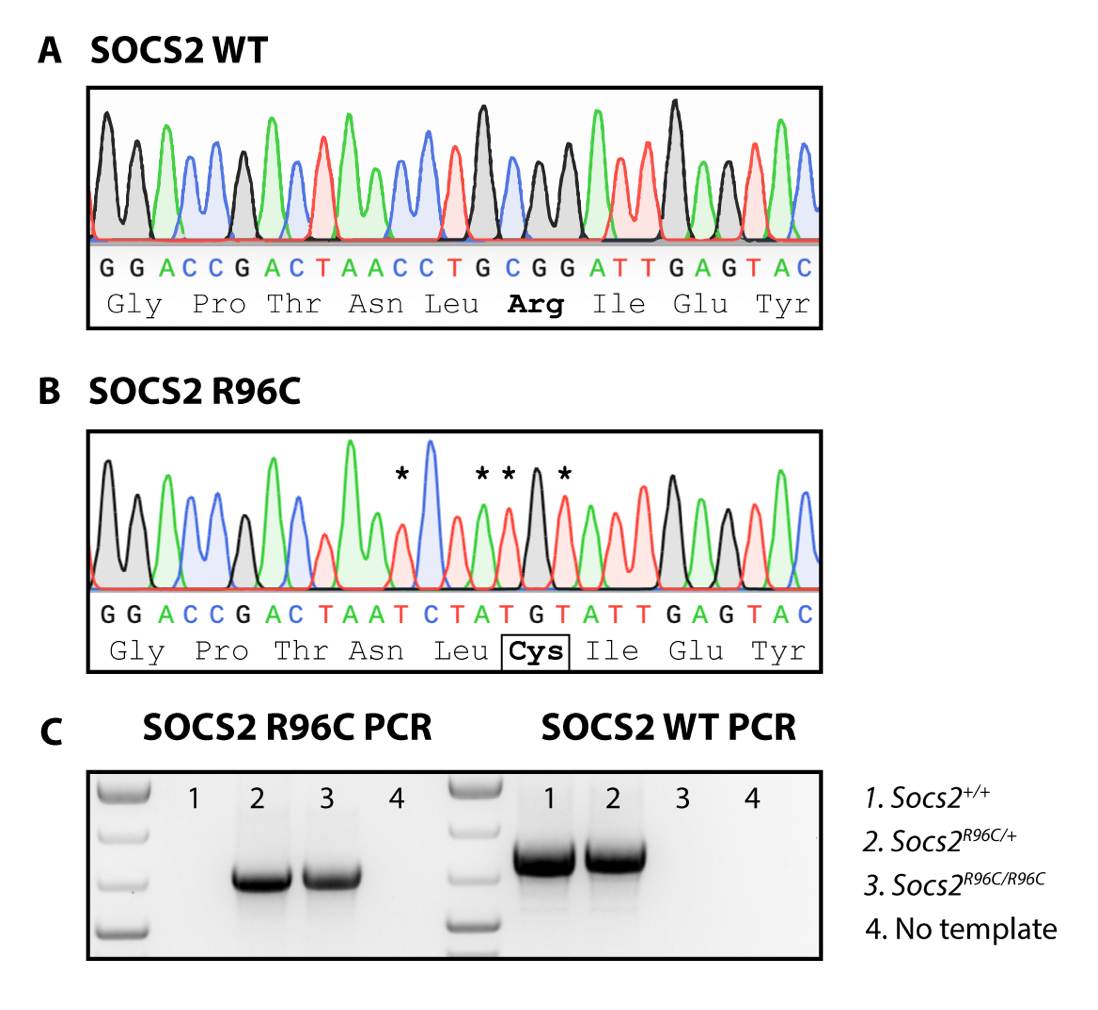

**Supplementary Figure 2. Genotyping confirmation of the correct sequence change in the *Socs2^R96C^* mouse.** Sanger sequencing chromatograms confirming **(A)** the WT *Socs2* sequence and **(B)** the mutated *Socs2^R96C^* sequence (homozygous mutant allele). *Highlights the CRISPR targeted base residue changes, two of which are synonymous and were introduced to enable primer specificity for standard gDNA genotyping. **(C)** Example 2% agarose gel confirming genotyping specificity by PCR. Primer sequences are available in **Supplementary Table 1**.

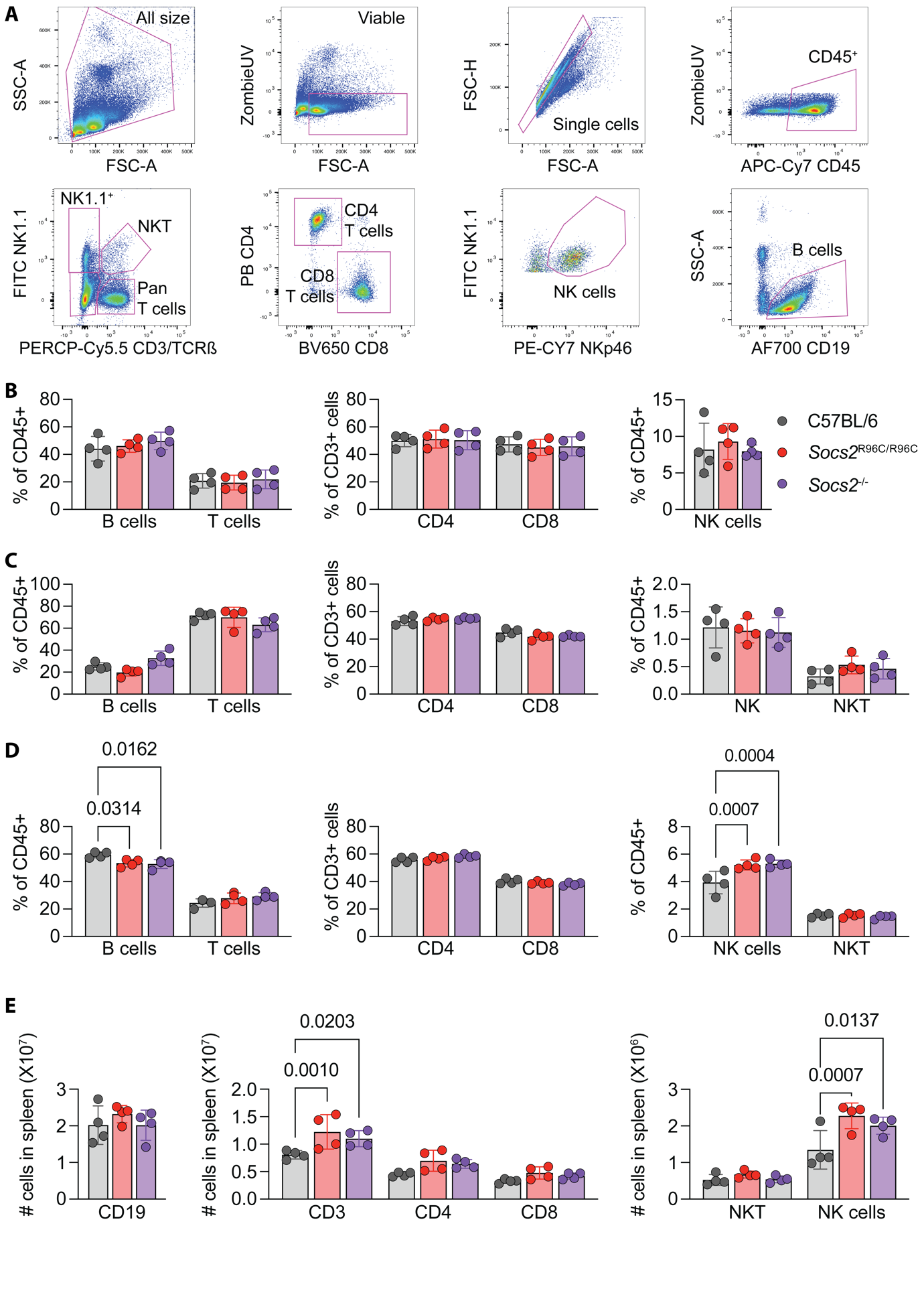

**Supplementary Figure 3. Homozygous *Socs2^R96C/R96C^* and *Socs2^-/^****^-^* **mice display increased numbers of splenic NK cells.** Single cell suspension peripheral blood mononuclear cells (PBMCs), splenocytes and lymphocytes from 8-12-week-old control C57BL/6 (black), *Socs2^R96C/R96C^* (red) and *Socs2^-/-^* (purple) mice were stained with fluorescently conjugated antibodies to various immune markers and analysed by flow cytometry. **(A)** Example gating for the analysis of viable leukocytes (ZombieUV^-^CD45^+^), NKT (CD3^+^TCRß^+^NK1.1^+^), Pan T cells (CD3^+^TCRß^+^NK1.1^-^), CD4 T cells (CD3^+^TCRß^+^NK1.1^-^CD4^+^), CD8 T cells (CD3^+^TCRß^+^NK1.1^-^CD8^+^), Pan B cells (CD3^-^TCRß^-^NK1.1^-^CD19^+^) and NK cells (CD3^-^TCRß^-^NK1.1^+^NKp46^+^). Percentage of various immune populations in the **(B)** blood, **(C)** axial and inguinal lymph nodes, and **(D)** spleen. **(E)** Enumeration of immune populations in the spleen using counting beads (123count eBeads). **(B-E)** Each dot represents cells from an individual mouse. Significance determined by two-way ANOVA with Sidak’s multiple comparisons test.

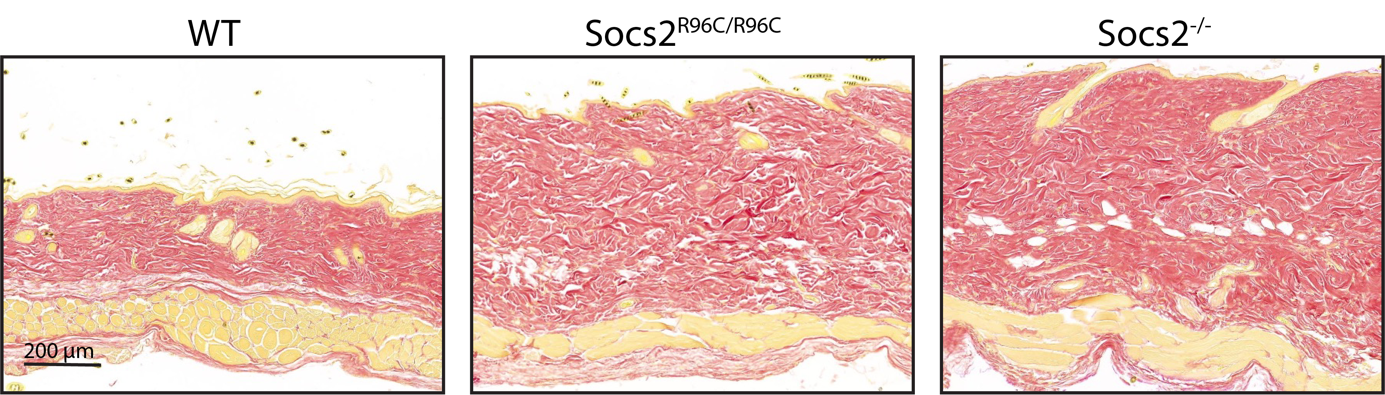

**Supplementary Figure 4. Homozygous *Socs2^R96C/R96C^* and *Socs2^-/^****^-^* **mice display thickening of the skin.** Van Giessen-stained dorsal skin section from 10-week-old male WT, *Socs2^R96C/R96C^* and *Socs2^-/-^* mice. *Socs2* mutant and null mice show increased collagen deposits and a thickened dermis. Representative images of n=3 mice/genotype. Scale bar = 200 µm.

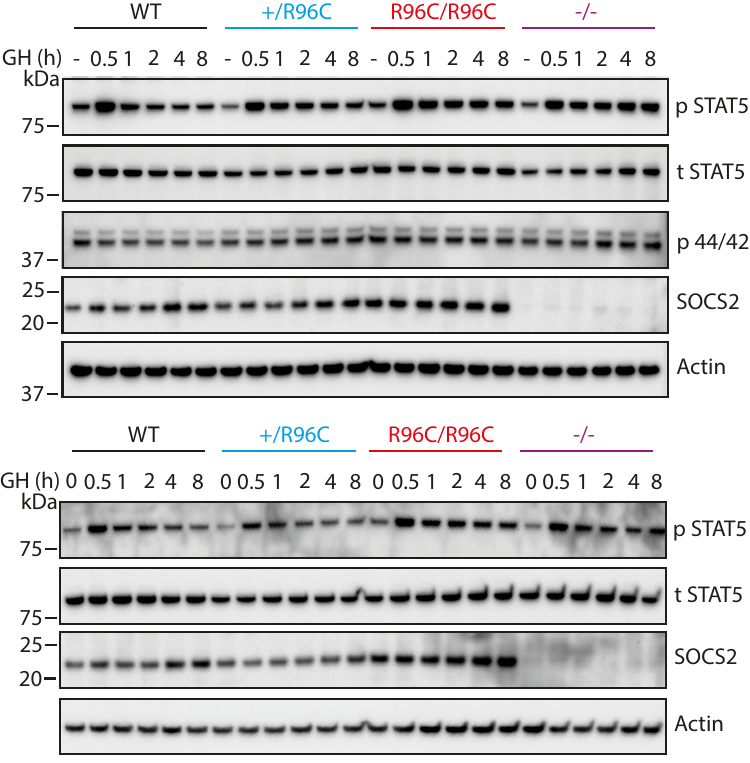

**Supplementary Figure 5.** ***Socs2^R96C/R96C^* and *Socs2^-/-^* MEFs display prolonged growth hormone signal activation.** *Socs2^+/+^*, *Socs2^R96C/+^*, *Socs2^R96C/R96C^* and *Socs2^-/-^* MEFs were treated with 50 ng/mL of GH, lysed and analysed by immunoblotting with antibodies to the indicated proteins. P: phosphorylated, T: total. Two additional independent experiments related to **Figure 4.**

**Supplementary Table 1. *Socs2^R96C^* genotyping primers.**

|  | **Allele** | **Forward Primer (5’-3’)** | **Reverse Primer (5’-3’)** | **PCR Product (bp)** |
| --- | --- | --- | --- | --- |
| **Standard genotyping** | *Socs2^R96C^* | AGCTTTCCACTTTGTCCCCTA | ATCTGAATTTCCCATCTTGGTACTCAATACAT | 766 |
|  | *Socs2^+/+^* | GCTGGACCGACTAACCTGC | AGCATGGTCAGCTTAACGGAA | 901 |
| **NGS* genotyping** |  | GTGACCTATGAACTCAGGAGTCTGACTGTTAATGAAGCCAAAGAG | CTGAGACTTGCACATCGCAGCGTGAACAGTCCCATTCCGTG | 290 |

*Next generation sequencing
